# Patterning defects in mice with defective ventricular wall maturation and cardiomyopathy

**DOI:** 10.1101/2025.05.22.655570

**Authors:** Javier Santos-Cantador, Marcos Siguero-Álvarez, José Luis de la Pompa

## Abstract

Ventricular chamber development involves the coordinated maturation of diverse cell populations. In the human fetal heart, single-cell RNA sequencing (scRNA-seq) and spatial transcriptomics reveal marked regional gene expression differences. In contrast, the mouse ventricular wall appears more homogeneous, except for a transient hybrid cardiomyocyte population co-expressing compact (*Hey2*) and trabecular (*Irx3, Nppa, Bmp10*) markers, indicating a transitional lineage state. To further investigate this, we used in situ hybridization (ISH) to examine the expression of a selected set of markers in normal and left ventricular non-compaction cardiomyopathy (LVNC) mouse models. In developing mouse ventricles, the expression of key marker genes was largely restricted to two wide myocardial domains— compact and trabecular myocardium—suggesting a less complex regional organization than human fetal heart. Other markers labelled endocardial and coronary endothelial cells rather than cardiomyocytes, differing from patterns observed in the human heart. In the LVNC model, various markers exhibited altered spatial expression, indicating that precise regional organization of gene expression is critical for normal ventricular wall maturation. These findings underscore the critical role of spatially regulated gene programs in ventricular chamber development and point to their potential involvement in cardiomyopathy pathogenesis.

## Introduction

The vertebrate heart starts as a linear tube and undergoes complex morphogenesis to form a four-chambered organ (1, 2). Central to this process is the coordinated growth and differentiation of the ventricular myocardium into trabecular and compact layers (3, 4). Trabeculation is the first sign of ventricular development, where cardiomyocyte projections lined by endocardium extend into the lumen, increasing surface area and supporting early cardiac output (5, 6). This is followed by compact myocardium expansion and trabecular remodeling, which contribute to ventricular wall thickening, the conduction system (7), and coronary vasculature (8). These processes result in a postnatal heart with a thick compact myocardium and smooth ventricular surface (9-11). Recent single-cell transcriptomic advances have revealed distinct gene expression profiles in myocardial regions (12-14). In developing human hearts, scRNA-seq combined with MERFISH (Multiplexed error-robust fluorescence *in situ* hybridization) has uncovered significant heterogeneity among ventricular cardiomyocytes (15). The trabecular myocardium is closely linked to the ventricular conduction system (VCS), which enables rapid electrical signaling (16). Markers like *IRX1–3, GJA5*, and *CGNL1* identify regions fated for VCS development (15). In contrast, the compact myocardium expresses *HEY2* and *PLK2*, linked to epicardial-proximal cardiomyocytes (15). Partial overlap of *IRX3* and *HEY2* defines a transitional hybrid zone (15), reflecting the spatial dynamics of ventricular maturation (10, 17). Genetically modified mouse models have shed light on pathways regulating ventricular patterning. NRG1 gain-of-function (*R26Nrg1^GOF^;Nkx2.5^Cre^*) causes premature VCS differentiation (18), while loss of MIB1-NOTCH signaling (*Mib1^flox^;Tnnt2^Cre^*) results in left ventricular non-compaction (LVNC), marked by excessive trabeculation, impaired ventricular wall maturation and cardiac dysfunction (9, 19). Temporally controlled inactivation of Nkx2-5 in trabeculae causes hypertrabeculation, fibrosis, and VCS hypoplasia (20). These models highlight the importance of precisely regulated gene expression in maintaining the structural and functional integrity of the ventricular wall.

In this study, we utilized ISH to map the expression of key ventricular wall markers in mouse hearts at embryonic day 15.5 (E15.5), corresponding to approximately 13 post-conception weeks (p.c.w.) in humans. This allowed cross-species comparison of spatial gene expression, revealing both conserved and species-specific features of ventricular wall maturation. We examined both normal and mutant mice to identify marker-associated defects. Our findings contribute to a molecular map of the developing ventricular wall, offering new insights into the genetic and cellular mechanisms underlying myocardial patterning and function.

## Materials and methods

### Mouse lines and genotyping

We used a conditional *Mib1^flox^*mouse line (9) and the myocardium-specific driver strain *Tnnt2^Cre^* (21). Both mouse lines were genotyped as described and maintained in a C57BL/6 inbred background. Animal studies were approved by the CNIC Animal Experimentation Ethics Committee and by the Community of Madrid (Ref. PROEX 054.6/25). All animal procedures conformed to EU Directive 2010/63EU and Recommendation 2007/526/EC regarding the protection of animals used for experimental and other scientific purposes, enacted in Spanish law under Real Decreto 118/2021 (modification on Real Decreto 53/2013) and Law 32/2007.

### Tissue processing

Mouse embryos were dissected at E15.5 in ice-cold PBS 1X and torsos were fixed overnight in 4% PFA at 4ºC using a shaker. After fixing, torsos were dehydrated through multiple graded ethanol washes and embedded in paraffin for microtome sectioning. Paraffin blocks were kept at room temperature prior to sectioning.

### Bright field imaging

Images of ISH stainings were obtained with an Olympus BX51 fluorescence microscope coupled to a Nikon DP71 camera and the CellSens Entry software (version 5.1).

### RNA-seq data analysis

Differential expression data on the *Mib1^flox^;Tnnt2^Cre^* (9) and *R26RNrg1^GOF^;Nkx2.5^Cre^* (18) models were obtained from previously published experiments. The analysis was done with the EdgeR R library, and differential gene expression was tested using a generalized linear model contained in the library. Genes showing altered expression with an FDR < 0.05 were considered differentially expressed.

### RNA probe synthesis and *in situ* hybridization on sections

Antisense RNA probes were designed to be exon spanning and complementary to the 3’ region of the transcripts of interest. Probes for *Irx3* (22), *Bmp10, Gja5, Anf* and *Hey2* (9) were described and used in previous literature. The remaining probes were generated using the primers listed below:

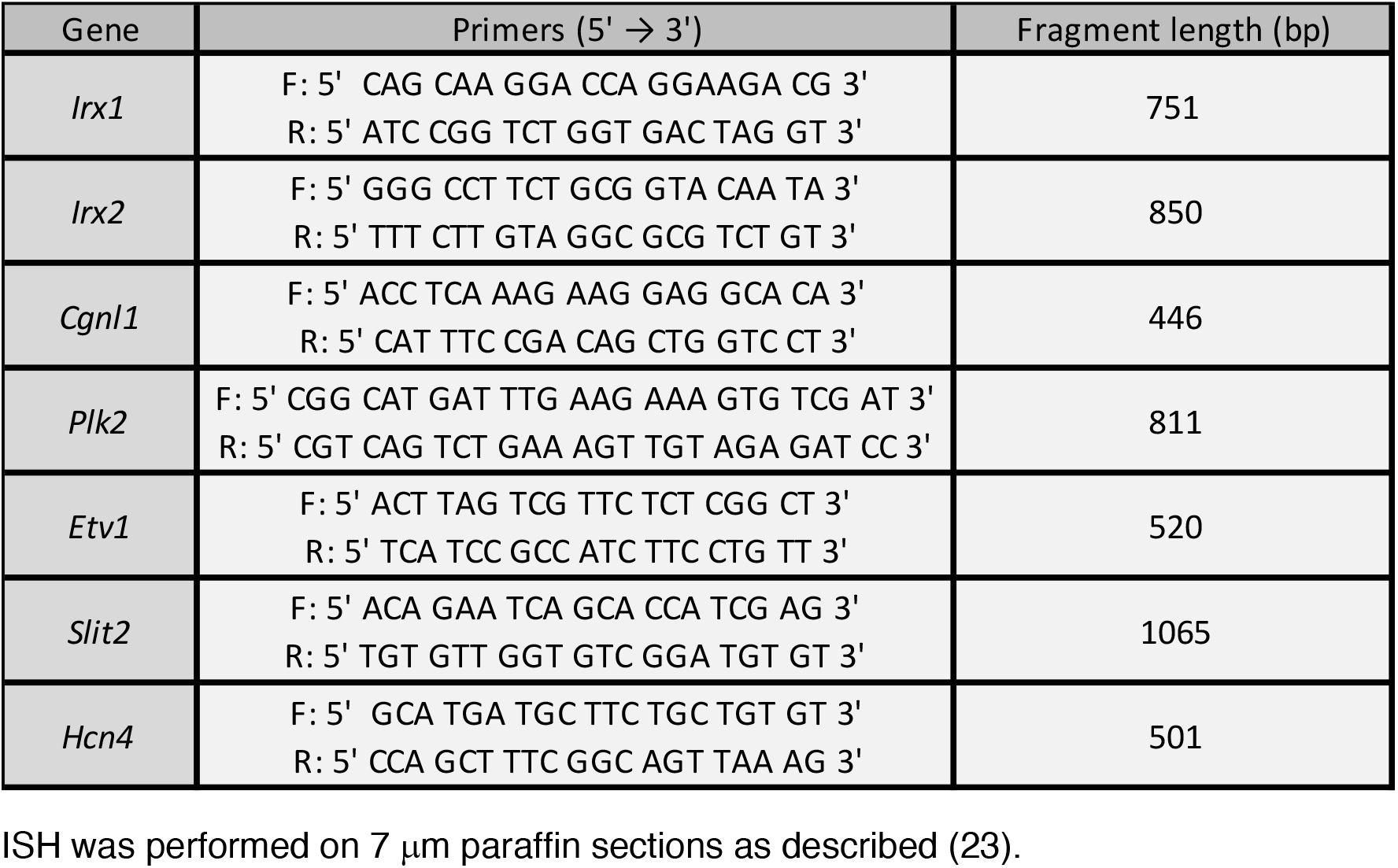

## Results

ScRNA-seq and MERFISH analyses of the developing human heart at 13 post-conception weeks (p.c.w.) have identified several genes differentially expressed across the ventricular wall (15). This stage is equivalent to embryonic day 15.5 (E15.5) in the mouse (**Figure 1A**). In the human heart, *IRX1* and *IRX2* are specifically expressed in the apical region of the trabeculae, within cardiomyocytes destined to form the VCS (15). The broader trabecular myocardium is characterized by enriched expression of *BMP10, IRX3, GJA5*, and *CGNL1* (**Figure 1B**). In contrast, the compact myocardium exhibits expression of *HEY2* and *PLK2*, with *PLK2* marking cardiomyocytes located closest to the epicardium (**Figure 1B**). Notably, the expression domains of *IRX3* in the trabecular myocardium and *HEY2* in the compact layer partially overlap in a transient region referred to as a “hybrid” ventricular myocardium region (15) (**Figure 1B**). This region, which marks the boundary between trabecular and compact myocardium, has been similarly identified in mouse models with impaired ventricular development (17) and during ventricular wall maturation (10). Within the compact layer, *PLK2* partially overlaps with *HEY2*, reinforcing its association with epicardial-proximal cardiomyocytes (**Figure 1B**).

**Figure 1.**
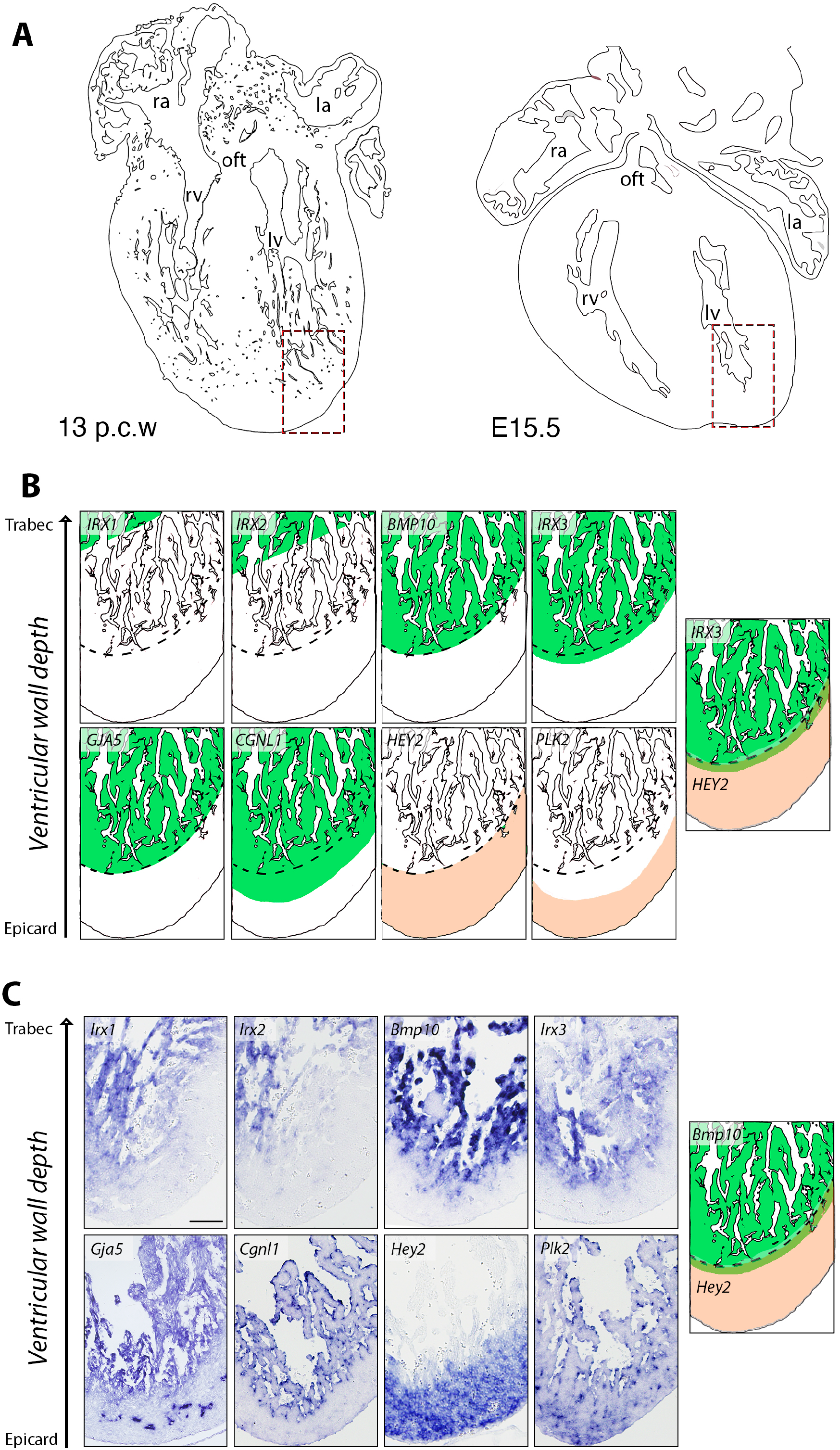
Comparative expression of marker genes in developing human and mouse ventricles. **(A)** Schematic representations of a 13 post-conception week (p.c.w.) human heart and an E15.5 mouse heart. **(B)** Schematic depiction of spatial gene expression patterns (*IRX1, IRX2, BMP10, IRX3, GJA5, CGNL1, HEY2* and *PLK2*) in the developing 13 p.c.w. human left ventricle, adapted from (15). Gene expression domains are shown in green and beige, for trabecular and compact myocardium genes, respectively. The scheme on the right shows the overlap between *IRX3* and *HEY2* expression domains. (**C**) ISH analysis of marker gene expression in the left ventricle of E15.5 wild-type mouse embryos. Note *Gja5* transcription in developing coronaries. The scheme shows the overlap between *Bmp10* and *Hey2* expression domains. Scale bar: 200 *μ*m.

We used ISH to examine the expression patterns of these marker genes in E15.5 wild-type mouse hearts (**Figure 1C**). *Irx1* and *Irx2* exhibited partially overlapping expression in the distal region of the trabecular myocardium, mirroring their spatial distribution in the developing human heart (15). *Bmp10, Irx3*, and *Gja5* were broadly expressed throughout the trabecular myocardium, with *Gja5* additionally marking the developing coronary vasculature. In contrast, *Hey2* was predominantly expressed in the compact myocardium (**Figure 1C**). Interestingly, the most basal/epicardial *Irx3* transcriptional domain partially overlaps with the most apical *Hey2* domain, defining a hybrid region of ventricular myocardium where *Irx3* and *Hey2* are co-expressed (15). A similar overlap is observed between *Bmp10* and *Hey2* expression domains (**Figure 1C**). Notable interspecies differences were observed for *Cgnl1* and *Plk2*: in the mouse heart, *Cgnl1* expression was weakly expressed in the myocardium whereas endocardial transcription was prominent (**Figure 1C**). *Plk2* was expressed in ventricular myocardium but was also detected in both the coronary endothelium and endocardium (**Figure 1C**). Shared domains of expression for genes such as *Irx1-3, Bmp10,* and *Hey2* support a conserved molecular architecture, including hybrid regions of co-expression that may reflect transitional myocardial zones. However, species-specific differences in *Cgnl1* and *Plk2* expression, particularly their enriched endocardial and endothelial localization in the mouse heart, underscore the importance of cross-species analysis for interpreting gene function and regulatory dynamics in ventricular development.

We next analysed the expression of these marker genes using RNA-seq data sets from two mouse models with altered ventricular development: one exhibiting premature differentiation of the ventricular conduction system (*R26Nrg1^GOF^*;*Nkx2.5^Cre^*) (18), and another displaying impaired ventricular wall maturation leading to cardiomyopathy (LVNC, *Mib1^flox^;Tnnt2^Cre^*) (9). In *R26Nrg1^GOF^*;*Nkx2.5^Cre^*hearts, we observed upregulation of markers associated with ventricular conduction system and trabecular myocardium differentiation, alongside downregulation of the compact myocardium marker *Hey2* (**Figure 2A**). In contrast, these same markers were altered in the opposite direction in *Mib1^flox^;Tnnt2^Cre^*hearts (**Figure 2B**). Together, these findings demonstrate that expression of ventricular wall markers is disrupted in both models, reflecting distinct yet complementary perturbations in myocardial patterning and maturation associated with disease.

**Figure 2.**
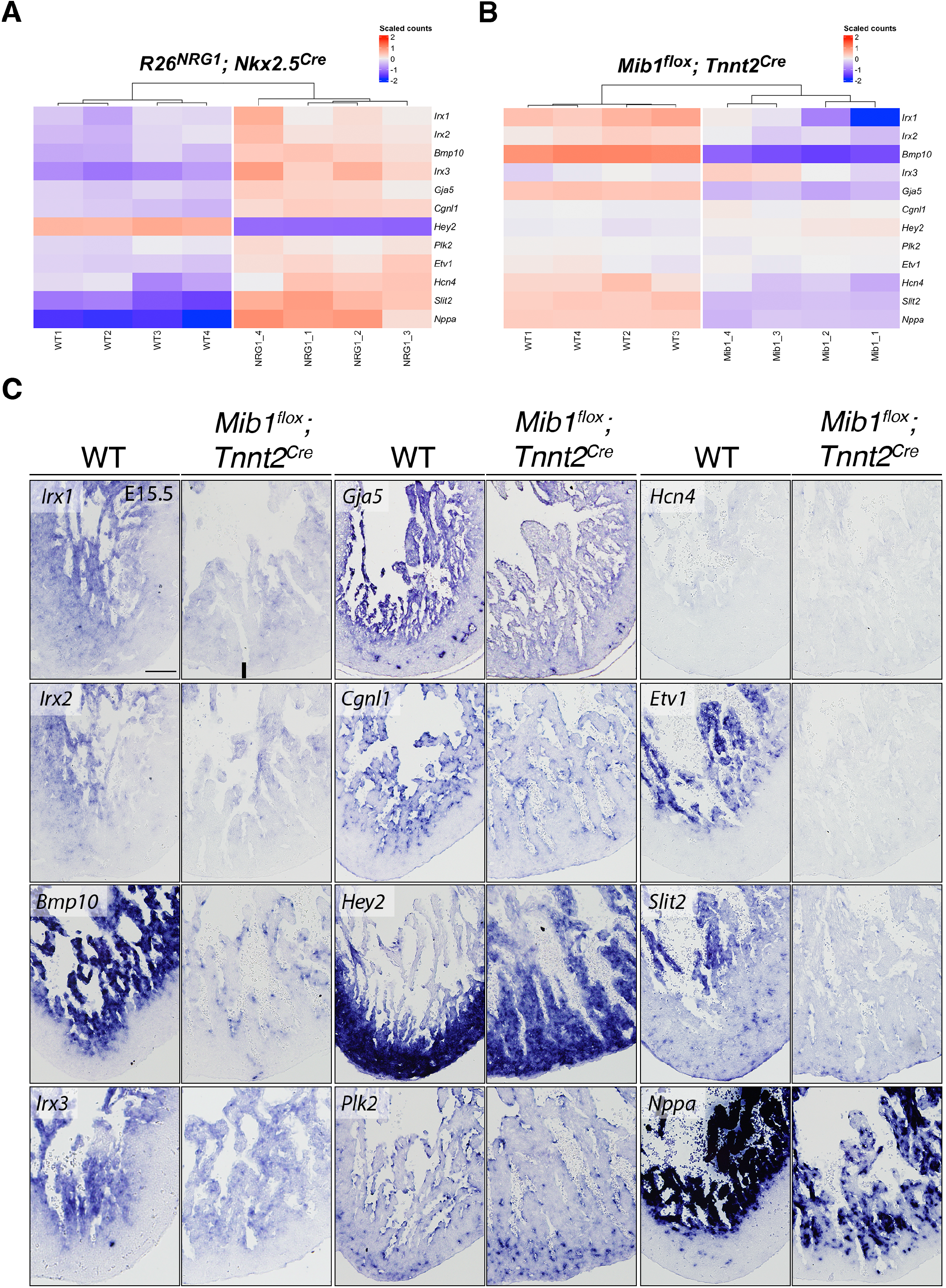
Dysregulated expression of ventricular wall marker genes in mouse models of conduction system mispatterning and cardiomyopathy. **(A)** Heatmap of differentially expressed genes (DEGs) in E15.5 wild-type (WT) and *R26Nrg1^GOF^*;*Nkx2.5^Cre^*hearts, highlighting upregulation of conduction system and trabecular markers, and downregulation of compact myocardium markers. **(B)** Heatmap of DEGs in E14.5 WT and *Mib1^flox^;Tnnt2^Cre^*hearts, showing reciprocal changes in gene expression relative to the *R26Nrg1^GOF^*;*Nkx2.5^Cre^*model. **(C)** ISH analysis of selected ventricular wall markers in E15.5 WT and *Mib1^flox^;Tnnt2^Cre^*hearts. The black bar indicates the thickness of compact myocardium in mutants. Scale bar: 200 *μ*m.

To directly compare with the LVNC phenotype, we performed ISH on heart sections from wild type (WT) and *Mib1^flox^;Tnnt2^Cre^*embryos at E15.5, using probes for additional trabecular myocardium marker genes, including *Hcn4, Etv1, Slit2*, and *Nppa* (**Figure 2C**). The observed expression patterns were largely consistent with our RNA-seq data, confirming altered expression of *Irx1, Irx2, Bmp10, Irx3, Gja5, Cgln1, Hey2, Hcn4, Etv1, Slit2* and *Nppa* in the mutant hearts (**Figure 2C**). Specifically, transcription of *Irx1-Irx3* was reduced in the trabeculae of *Mib1^flox^;Tnnt2^Cre^*hearts, although their spatial localization remained largely unchanged. In contrast, *Bmp10, Gja5, Etv1, Slit2* and *Nppa* showed markedly reduced expression in trabecular myocardium, indicating a more pronounced disruption of trabecular gene programs (**Figure 2C**). *Cgln1* expression was specifically diminished in the endocardium while *Plk2* appeared unaffected (**Figure 2C**). Notably, *Hcn4* expression was slightly reduced in mutants (**Figure 2C**), consistent with its naturally low levels in the ventricular myocardium at later developmental stages, when the ventricular conduction system becomes more specialized (24). These findings suggest that impaired maturation of the trabecular myocardium in *Mib1^flox^;Tnnt2^Cre^*mutants disrupts conduction system development, potentially contributing to the pathophysiology of LVNC.

The compact myocardium marker *Hey2* exhibited an expanded expression domain extending into the trabeculae in *Mib1^flox^;Tnnt2^Cre^*hearts (**Figure 2C**). This agreed with a mild *Hey2* upregulation observed in the RNA-seq of these mutants (**Figure 2B**) and (9). This suggests a disruption in the spatial patterning of the ventricular wall in the context of *Mib1* loss of function. Together, these results confirm and extend our RNA-seq findings, supporting a model in which *Mib1* deficiency disrupts the coordinated gene expression programs necessary for normal spatial organization and maturation of the ventricular wall.

## Discussion

In this study, we analyzed the expression of markers identified in the 13 p.c.w. human ventricular wall (15) in the E15.5 mouse heart, two developmentally equivalent stages. Our findings reveal that these markers define distinct regions within the trabecular myocardium or span the entire compact myocardium, reflecting a more streamlined organization of the developing mouse ventricles. Notably, the trabecular myocardium can be further subdivided into apical cardiomyocytes, which contribute to the formation of specialized cells within the VCS (16), and basal trabecular cardiomyocytes (6).

Recent work by Cui et al. using snRNA-seq and spatial transcriptomics in E15.5 mouse hearts, identified seven ventricular cardiomyocyte populations classified according to marker expression levels and spatial localization (25). Comparison with the eight populations identified by scRNA-seq and MERFISH in human hearts (15), revealed overlapping genes expression signatures in compact (*Hey2*) and trabecular myocardium (*Bmp10, Irx1,2, Gja5*), as confirmed by our ISH results. These findings highlight conserved gene expression signatures despite differences in overall complexity between mice and humans.

Farah et al. (2024) identified in the developing human heart a transient hybrid ventricular cardiomyocyte population co-expressing the trabecular marker *IRX3* and the compact marker *HEY2* (15). This aligns with earlier lineage-tracing studies by Tian et al. (2017), which identified a hybrid myocardial zone in early postnatal mouse ventricles, formed by cells originating from both *Nppa*^*+*^(trabecular) and *Hey2*^*+*^(compact) lineages (10). Feng et al. (2022) also identified transient hybrid cardiomyocytes during specific developmental windows in mice (26). These data reveal the conservation between humans and mice of a transient hybrid ventricular cardiomyocyte population that likely represents the developmental transition of cardiomyocytes originated in the compact myocardium towards a trabecular fate, with the progressive downregulation of compact markers (*Hey2*) and upregulation of trabecular ones (*Irx3, Nppa, Bmp10*). Notably, studies in mice harbouring inactivating mutations in the NOTCH signalling regulator Mindbomb1 (Mib1) or the ligands Jag1 and Jag2 revealed a transient “intermediate myocardium” zone within abnormal trabeculae, composed of *Hey2^+^, Bmp10-, Cx40*-cardiomyocytes, which progressively develop a left ventricular non-compaction (LVNC) phenotype (9, 17, 19). Interestingly, we previously identified a distinct inner compact myocardium region adjacent to the trabecular myocardium that is absent in mice with endothelial *Nrg1* loss- or myocardial Nrg1 gain-of-function (18), highlighting the sensitivity of the compact-trabecular transition zone to key signalling pathways. These findings deepen our understanding of ventricular wall morphogenesis and underscore the importance of this transitional region in the pathogenesis of congenital heart disease and cardiomyopathies.

Several genes with altered expression, including *Irx1-3, Gja5*, and *Etv1,* are implicated in VCS development and function. Given that trabeculae are the progenitors of the VCS (27), dysregulation of these markers may contribute to the arrhythmias and conduction defects seen in LVNC. *Irx1* loss-of-function in zebrafish leads to a dose-dependent heart rate reduction of up to 65% (28, 29), while *Irx2* appears largely redundant. In contrast, Irx3 acts as a key regulator of gap junction proteins, promoting *Gja1/Cx43* and suppressing *Gja5/Cx40* expression (30). Although *Irx3* shows minor expression changes in *Mib1^flox^;Tnnt2^Cre^*mutant mice, the downregulation of *Gja5* suggests additional regulatory mechanisms may contribute to the arrhythmogenic phenotype associated with LVNC. Etv1 is a conserved marker of trabecular myocardium in embryos and the VCS in postnatal hearts, in both mice and humans (15, 26). Its expression is positively regulated by Nrg1 via the Ras-MAPK pathway, as supported by our RNA-seq data and previous studies (31). *Etv1* inactivation in mouse models leads to VCS hypoplasia and reduced expression of *Gja5* and other conduction genes, resulting in defective conduction (31). Furthermore, human *ETV1* variants have been associated with conduction disease (31), highlighting the potential role of the Nrg1-Etv1 axis in the pathogenesis of LVNC and related cardiomyopathies.

Although *CGNL1* and *PLK2* show prominent expression in cardiomyocytes in human hearts (15), they are only weakly expressed in cardiomyocytes of developing mouse hearts. Instead, ISH and scRNA-seq data indicate that their transcription is primarily restricted to endocardial and endothelial cells (26, 32, 33). *Plk2* is also detected in fibroblasts, where it plays a critical role in maintaining normal cardiac function, as loss-of-function models exhibit pro-fibrotic and arrhythmogenic phenotypes (34, 35). While these markers are valuable for tracing endothelial lineages, their differential expression between mouse and human hearts remains an open question, challenging assumptions about conserved developmental pathways.

In conclusion, our findings show that while the developing mouse and human hearts share conserved gene expression signatures and transitional cardiomyocyte populations, key differences in gene expression levels, cellular composition, and regulatory dynamics persist between species. The identification of hybrid and intermediate myocardial zones in both humans and mice emphasizes the importance of transitional states in ventricular wall maturation and highlights their vulnerability to disruption by genetic or signalling perturbations. Moreover, the distinct expression profiles of endothelial-enriched genes such as *Cgnl1* and *Plk2* point to species-specific adaptations in cardiac development. Continued cross-species analyses integrating spatial, transcriptional, and functional data will be critical for refining our understanding of ventricular morphogenesis and for improving the translational relevance of mouse models in the study of congenital heart disease and cardiomyopathies such as LVNC.

## Author Contributions

J.S.-C., M.S.-A., and J.L.d.l.P. conceptualized the study and design of methodology. J.S.-C. and M.S.-A. carried out experiments and acquired images. J.S.-C., M.S.-A., and J.L.d.l.P. analyzed and interpreted data. J.S.-C., M.S.-A., and J.L.d.l.P. wrote the manuscript. J.L.d.l.P. provided funding and material support. All authors have read and agreed to the published version of the manuscript.

## Funding

This study was supported by grants PID2022-136942OB-I00 and CB16/11/00399 (CIBER CV) from MICIU/AEI/10.13039/501100011033, and grants for Research in Genetic Cardiomyopathies (SECSCFG-INV-CFG 21) from the Spanish Society of Cardiology to J.L.d.l.P. The cost of this publication was supported in part with funds from the European Regional Development Fund. The CNIC is supported by the ISCIII, the MCIN and the Pro CNIC Foundation and is a Severo Ochoa Center of Excellence (grant CEX2020-001041-S) financed by MCIN/AEI /10.13039/501100011033.

## Institutional Review Board Statement

Animal studies were approved by the CNIC Animal Experimentation Ethics Committee and by the Community of Madrid (Ref. PROEX 054.6/25). All animal procedures conformed to EU Directive 2010/63EU and Recommendation 2007/526/EC regarding the protection of animals used for experimental and other scientific purposes, enacted in Spanish law under Real Decreto 118/2021 (modification on Real Decreto 53/2013) and Law 32/2007.

## Acknowledgements

We thank D. MacGrogan for critical reading of the manuscript, and A. Galicia and L. Méndez-Peralta for mouse husbandry.

## Conflicts of Interest

The authors declare no conflicts of interest.

## Notes

### Competing Interest Statement

The authors have declared no competing interest.

